# Efficient Techniques for Comprehensive Tissue Sampling in Adult Xenopus

**DOI:** 10.1101/2025.01.27.635174

**Authors:** Rachael A Jonas-Closs, Cora E Anderson, Leonid Peshkin

## Abstract

*Xenopus* has long been a pivotal model organism for investigating vertebrate development and disease, offering deep insights into cellular processes and gene function. Despite the wealth of information on embryonic *Xenopu*,*s* there remains a significant gap in standardized methods for adult tissue sampling, especially for modern approaches like quantitative proteomics. This study introduces a comprehensive protocol for rapid, precise, and efficient sampling of multiple tissues in adult *Xenopu*.*s*The protocol addresses challenges associated with the subtle anatomical differences compared to other anurans, ensuring reproducibility even for those with limited experience in frog dissection. This protocol is optimized for high-quality biochemical analyses by prioritizing sample freshness. We are facilitating the rapid collection of up to 18 tissues within an hour. Additionally, the methods apply to perfused and unperfused conditions, providing flexibility for a range of experimental needs. This work not only fills a critical methodological gap for *Xenopus laevis*and *tropicalis*but also serves as a valuable resource for researchers adapting techniques to similar amphibian models, thereby enhancing the scope and reliability of comparative biological and evolutionary studies.

**Summary:** This is part one of a comprehensive *Xenopus* sampling protocol. The tissues sampled are the heart ventricle, arterial trunk, left liver lobe, gallbladder, lung, pancreas, spleen, larynx, esophagus, stomach, intestines, testes, fat bodies, oviduct, paired kidneys, sciatic plexus, skin, thymus, and whole eye.

## Introduction

This paper aims to offer clear guidance for precise and reproducible organ sampling of adult *Xenopus* tissues, as is available for *Xenopus* tadpoles^1^. While a “digital dissection” of adult *Xenopus* is available^2^, achieving consistent organ and tissue sampling in adult *Xenopus* remains difficult without the detailed instructions provided for other adult models, such as mice^3,4,5^.

Although a comprehensive dissection guide exists for *Rana*sp.^6^ and various classroom dissection manuals are available for other anurans^7^, no specific dissection and sampling guide for *Xenopus* is currently available. For individuals who are inexperienced in sampling techniques or amphibian anatomy, the subtle differences between *Xenopus* and other anurans make existing resources inadequate for consistent and replicable tissue sampling.

This guide was developed to prioritize tissue freshness for proteomics and immunochemistry. To an experienced user, all tissues can be collected from one individual in under an hour. It is recommended that if user lacks experience they attempt this protocol on specimens that were euthanized for other reasons before sacrificing any animal that is more challenging to replace.

### PROTOCOL

**All experiments were performed in accordance with the rules and regulations of the Harvard Medical School IACUC (Institutional Animal Care and Use Committee). (IS 00001365_3).**

If perfusion protocol **8** is being followed prior to sampling skip to 2:2 (step 2 of sampling A) Preparation (without perfusion)

1. Ensure that the research institution has approved the euthanasia and perfusion technique described in this protocol.
2. Prepare a solution of 5g/L MS-222 (tricaine methanesulfonate) and 5g/L sodium bicarbonate (see **Table of Materials**). The volume should be greater than the volume required to completely cover the animals being euthanized. Check the pH to ensure that it is ≥7.
3. Perform primary euthanasia by placing the *Xenopus* in this solution (from step 2), the animal will remain submerged for a total of one hour.
4. Place the dissection surface (tray or foam sheet) at an incline within a secondary container or otherwise arrange it to facilitate blood drainage.
5. Once the frog has been in the solution for 1 hour, primary euthanasia has been completed. Remove the frog and check the loss of pain response by performing a foot pinch.
6. Record the age (if known), sex, and health status of the animal you are sampling, as well as if it was perfused. Weigh the Xenopus and take any additional measurements required before sampling.
7. Place the frog on its back and pin down the forelimbs proximal to the body (figure 1).
8. Using dissection scissors cut through the skin, up the midline, and then laterally, making two flaps.
9. Identify the linea alba (figure 2). Use forceps to grasp the linea alba and pull it away from the coelomic cavity. Carefully use scissors to cut up through the musculature. Make two flaps out of the cavity wall and cut or pin all flaps out of the way.
10. Identify the heart, which should still be beating. If the heart has stopped beating prior to sampling it should be noted that the sample freshness has been compromised. If the heart is not easily accessible, use dissecting scissors to reduce the coracoid bones (figure 2).
11. If desired complete the rapid perfusion protocol.
12. B) **Sampling**

**Figure 1:**
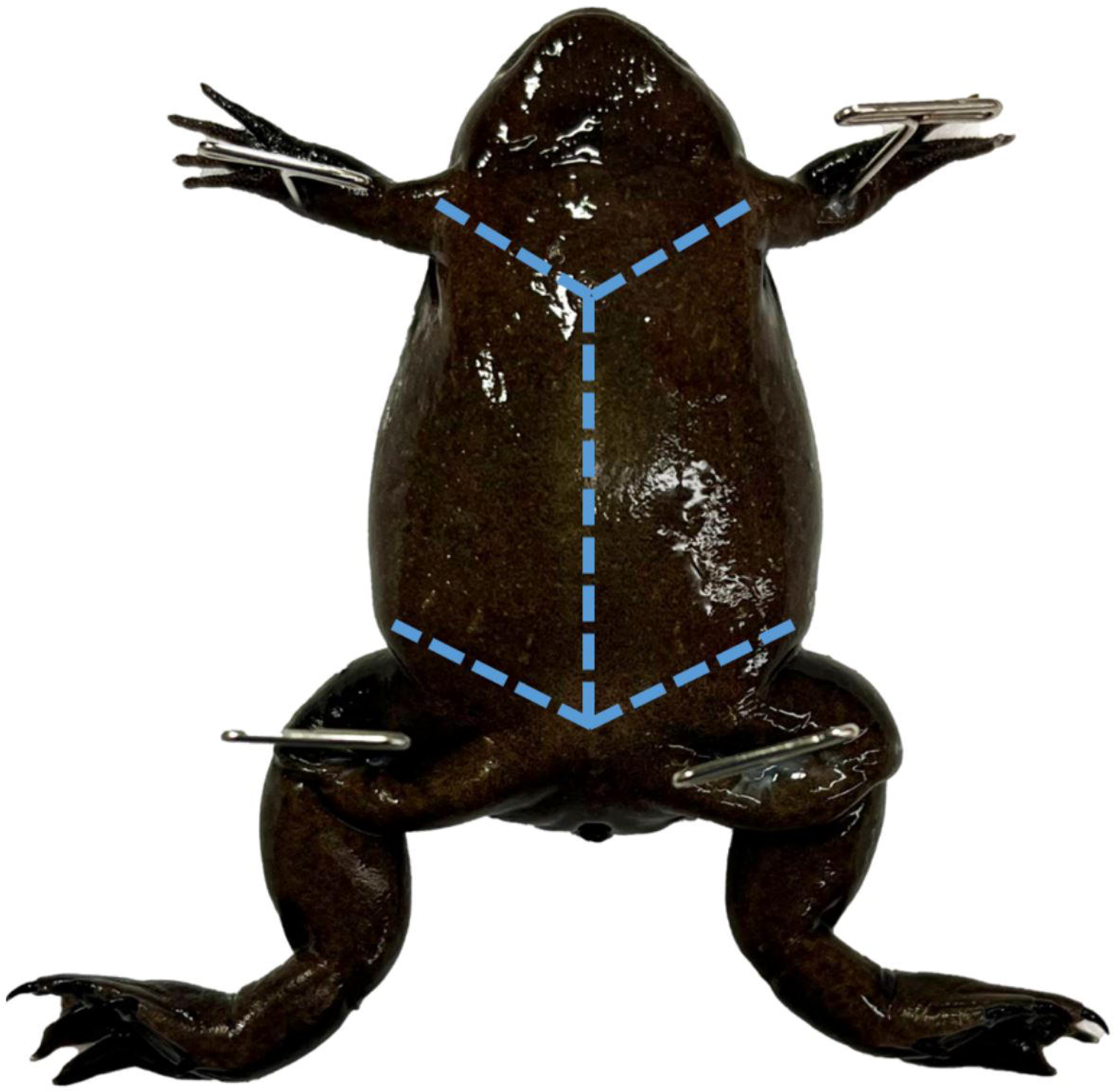
Pinned *Xenopus*. A mature female *X. tropicalis*pinned through each limb. Blue dashed lines indicate where the skin should be cut to generate two flaps.

**Figure 2:**
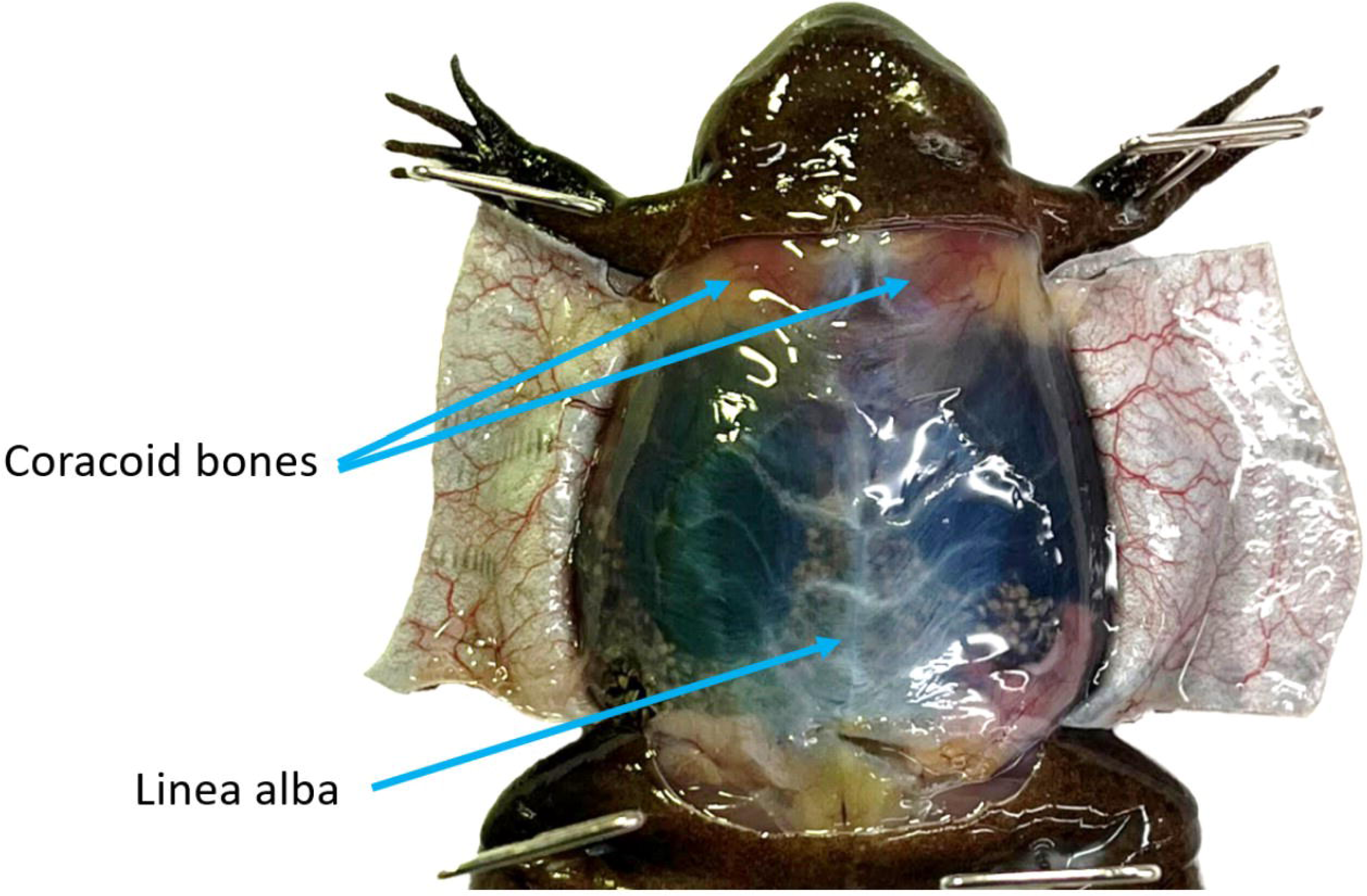
Abdominal wall. The ventral skin of an *X. tropicalis*female is cut into flaps, making the linea alba and coracoid bones visible.

**If the animal has been perfused, skip to step 3.**

**Note: all margins should be approximately 25mm. Forceps and scissors should be wiped clean or replaced between samples.**

All tissues should be sampled within 1 hour of euthanasia.

For unperfused animals, the coelomic cavity should be rinsed with chilled PBS or .7xPBS to clear blood and maintain visibility.

Visually inspect samples under magnification to ensure no unwanted tissues are attached.

Immediately after sampling tissues should be rinsed in chilled PBS or .7xPBS, depending on experimental needs^9^. In perfused animals, it is expected that all tissues excluding the liver will rinse cleanly. If this is not the case then a note should be made. In unperfused tissues, all rinsing media will be saturated with blood.

Use a P200/P20 to remove as much of the remaining buffer as possible, following this the samples should be flash-frozen in liquid nitrogen.

1. Identify the thin pericardium (figure 3) and pull it taut with tissue forceps. Using the tip of the iridectomy scissors gently perforate it, careful not to cut the underlying tissues. Peel the pericardium up away from the 3 chambers of the heart (figure 4a).
2. Trim away the auricles by cutting them where they meet the ventricle and arterial trunk.
3. Use forceps to grasp the ventricle by the apex and cut it below where it meets the arterial trunk, leaving a margin (figure 4b). Note that the center of the ventricle may include light-colored spiral valve tissue, this may be kept. In unperfused animals, the removal of the ventricle may qualify as secondary euthanasia.
4. Observe that the arterial trunk bifurcates into two aortic trunks (also called lateral aortae) and that each splits into three arterial arches (figure 4c). Pull the arterial trunk taut and cut the aortic trunks immediately after where these arches split.
5. The 3 lobes of the liver will be visible (figure 5). Grasp the lip of the left lobe (on the viewer’s right) and gently lift it so that the hepatic and cystic ducts are visible. Sample the bottom 2/3 of the lobe, below these attachments (figure 6a).
6. Identify the gallbladder, which may vary drastically in color. Grasp the gallbladder and pull it away from the right lobe of the liver, sever its attachment (figure 6b). If it does not burst then express it with iridectomy scissors.
7. Grasp the apex of the lung and pull it taut. Note the texture of the alveoli, and determine where that texture ends and the bronchus begins. Cut at this margin (figure 6c). The lung may be fused to the peritoneum, if this is the case, carefully detach it.
8. Removing the ovary is helpful to gain better access to a female frog’s tissues. Identify the ovary which is enveloped in a layer of visceral peritoneum called the germinal epithelium. Gently shift the lobes until they are on their respective sides to make the area of attachment visible (figure 7a). These attachments are directly ventral to the paired kidney. Using scissors, remove the ovaries as close to the kidneys as possible, without damaging them (figure 7b). During this process oocytes may leak out, these will be rinsed off after sampling.
9. Inspect the anterior lobe (also called the median lobe) of the liver and note how it connects to the stomach and duodenum through the mesentery (figures 5 and 8a). Server the mesentery and hepatoduodenal ligament using iridectomy scissors. The remaining fleshy mass is the pancreas and common bile duct.
10. Sever the connection of the pancreas and common bile duct to the anterior lobe of the liver, leaving a margin. Grasp the stomach with toothed forceps and one end of the pancreas with tissue forceps. Under magnification gently tease the pancreas off of the stomach (figure 8b). Should it not come away cleanly, the remaining pancreatic tissue will be visible and can be picked off. Alternatively, the pancreas can be methodically detached using iridectomy scissors and tissue forceps. In addition to the common bile duct, it is expected that some mesentery and the pancreatic duct will be included in this sample.
11. Identify the spleen and sample it by cutting its attachment to the peritoneum (figure 8c).
12. Remove and dispose of the remaining liver tissues.
13. Remove the other lung and the remaining bronchus where they attach to the larynx. Grasping the larynx by its inferior end, begin to lift it out of the cavity, severing clear attachments as they become apparent.
14. As the larynx is lifted its attachment to the esophagus will become noticeable. Carefully peel and cut the esophageal tissue away from the larynx using curved iridectomy scissors (figure 9). Then pull the larynx down and trim it away from the mouth cavity. Clear attachments will become apparent and should be cut.
15. Notice the curvature of the stomach which differentiates it from the esophagus and the pyloric restriction that differentiates it from the intestines(figure 10). Cut the alimentary canal immediately anterior to the stomach’s curvature, separating it from the esophagus.
16. Open the mouth and identify the shared opening of the eustachian tubes on its roof^10^. With the mouth open tug on the esophagus to note its attachment (figure 11a). Lift the esophagus out of the cavity severing clear attachments as you go. Sever the esophagus from the buccal cavity immediately below the eustachian opening.
17. Grasp the stomach and cut it where it meets the duodenum, at the pyloric restriction (figure 10).
18. Pull the colon taut and cut it as close to the cloaca as possible (figure 11b). Pull the intestines out, severing any clear peritoneal attachments.
19. If the frog is male identify the testes and remove one, using iridectomy scissors, being careful not to damage the underlying kidney (figure 12a).
20. Tease apart the fat bodies so that they are on their respective sides, the area, over the kidney, where the fat body connects to the peritoneum will be visible. Grasp the base of either fat body and use scissors to cut it away from the peritoneum, leaving a margin (figure 12b).
21. Identify and remove the urinary bladder, cutting as close to the cloaca as possible (figure 13a).
22. If the frog is female or has a distinct vestigial oviduct, grasp the inferior end of the oviduct, pull it away from the cloaca, and cut it as close to the cloaca as possible (figure 13b). Pull the oviduct out of the coelomic cavity, severing any peritoneal attachments as they become apparent (figure 13c).
23. Remove and dispose of the remaining fat bodies and testes or oviduct.
24. Use forceps to grasp the kidneys at their inferior base and note that they are covered by the clear peritoneum (retroperitoneal)^11^. Sever the peritoneum at the base of the kidney (figure 14a) and then lift the kidneys out of the coelomic cavity (figure 14b), using scissors to sever the peritoneum as needed.
25. Under magnification, cut away excess peritoneum and any remaining fat body, ovary, or testes tissue.
26. Note the bundle of peripheral nerves attaching the terminus of the spinal column to each leg, this is the sciatic plexus. Grasp the bundle of nerves with forceps and sever them where they exit the coelomic cavity (figure 14c). Lift the bundle out of the cavity and cut it where the nerves leave the vertebral column (figure 14d).
27. Remove the pins from the animal, flip it onto its ventrum, and re-pin the animal’s limbs. Select either hindlimb to sample from and pin the foot of that limb.
28. Remove an almond-shaped flap of skin from over the gastrocnemius/tibiofibula (figure 15).
29. Grasp the skin of the dorsum and cut laterally so there is a continuous cut around the midsection of the animal (figure 16a).
30. Remove the pins, secure the legs in one hand, and grasp the anterior dorsal skin with toothed forceps. Skin the animal by pulling the dorsal skin up over the head and arms, using force (figure 16b).
31. Identify the fatty mass covering the jaw joint. Sever the posterior margin of the mass from where it is fused to pink muscle tissues and then use iridectomy scissors to cut it away from the underlying bone (figure 17a). Depending on the animal’s age and maturity, a cluster of melanophores will become apparent (figure 17b). Trim the mass down so that no muscular attachments or cartilage are attached.
32. Repin the animal and select an eye. Insert curved iridectomy scissors into the orbit around the curvature of the eye (figure 17c). Gently cut around the eye, severing muscular attachments.
33. Slide the scissors deeper into the orbit and detach the optic nerve, behind the eye. Use these scissors to dislodge the eye out of the orbit and then further reduce any muscular attachments.

**Figure 3:**
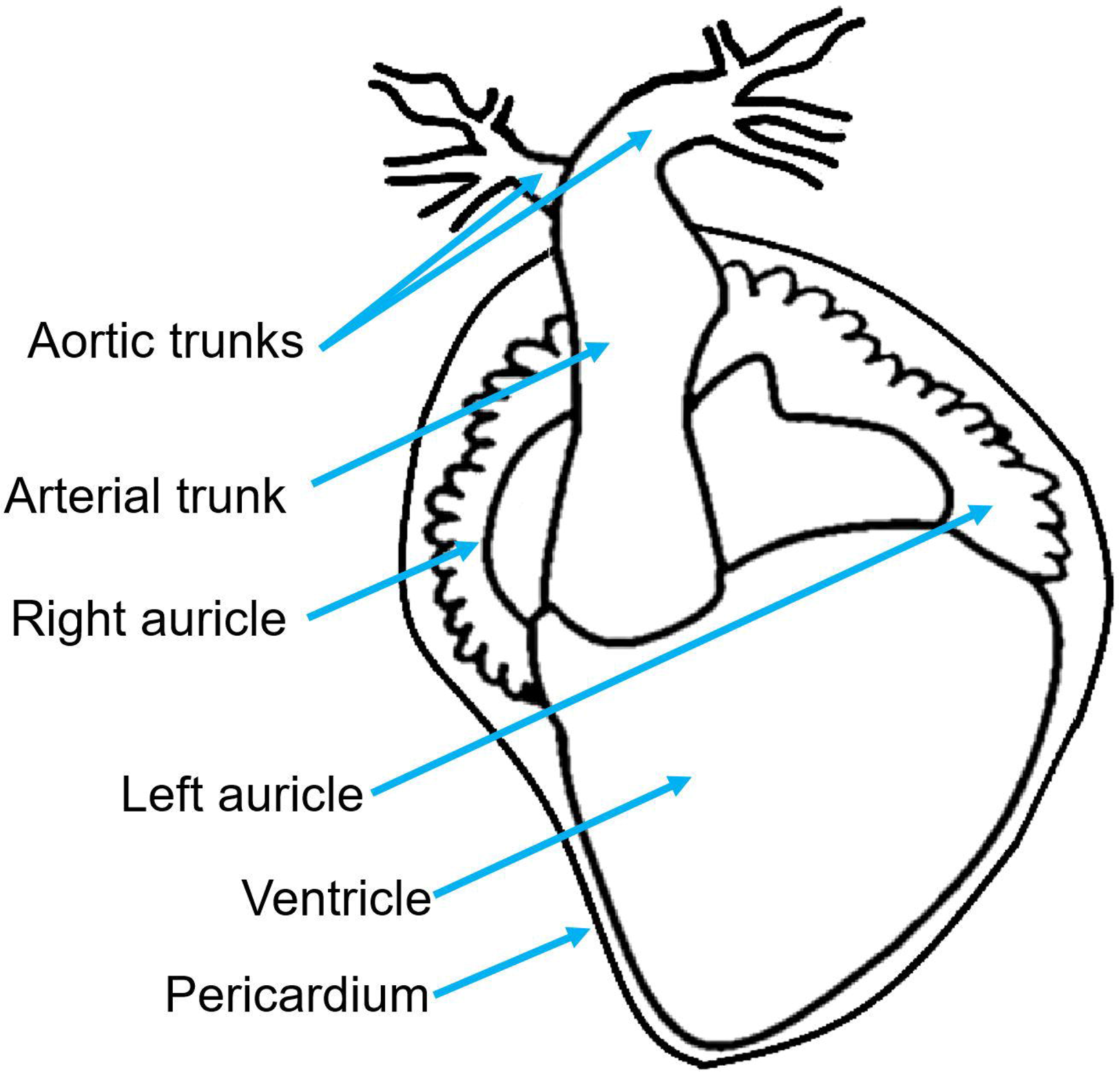
Xenopus heart diagram. A diagram of the Xenopus heart including the pericardium, ventricle, auricles, arterial trunk, and aortic trunks with their arches.

**Figure 4:**
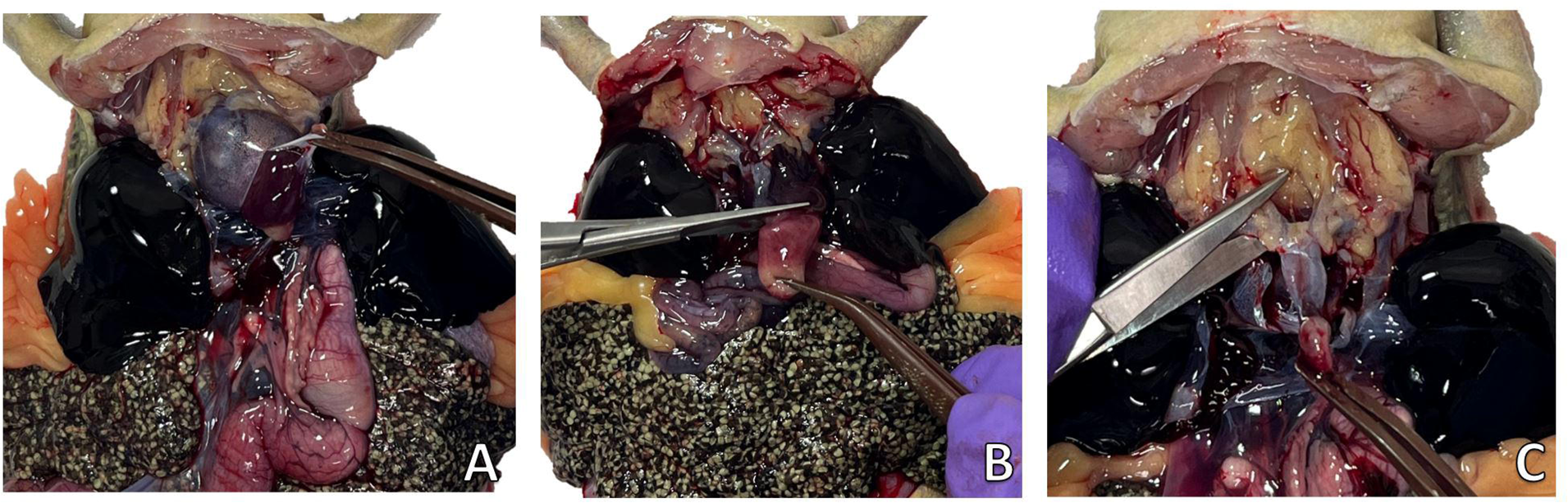
Ventricle and arterial trunk sampling of an unperfused *X. laevis* . (**A**) The pericardium being peeled up away from the heart, to reveal the three chambers. (**B**) The ventricle being sampled immediately below where it meets the atria and arterial trunk. (**C**) The arterial trunk and aortic trunks are disconnected from the arterial arches directly after their split.

**Figure 5:**
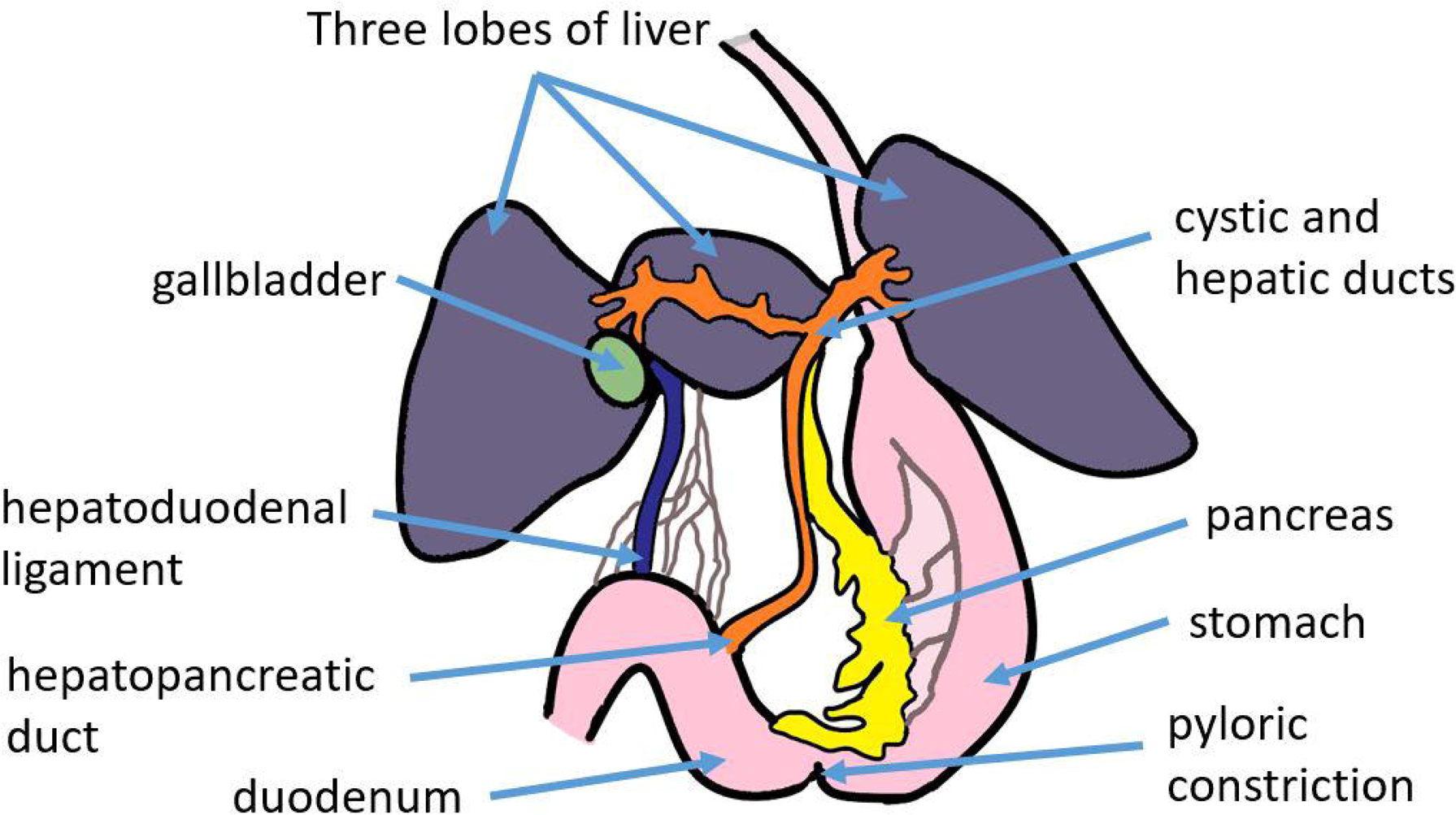
Hepatopancreatic diagram. A diagram of the 3 lobes of the liver, pancreas, and associated organs.

**Figure 6:**
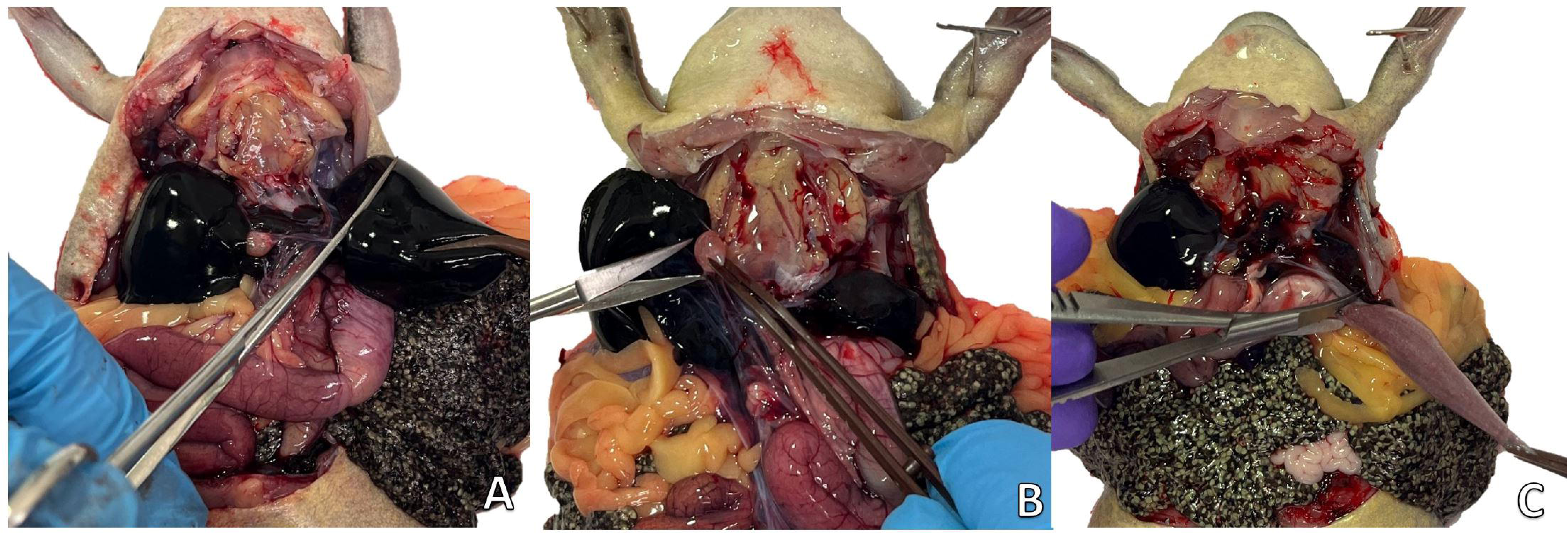
Liver, gallbladder, and lung sampling of an unperfused *X. laevis* . (**A**) The left liver lobe is pulled to the side, making its attachments to the hepatopancreatic ducts apparent. The liver is being sampled below these attachments. (**B**) The gallbladder being sampled. The gallbladder can vary highly in color depending on its bile content. (**C**) The lung is pulled down so that the distinction between it and the bronchus is visible and sampled at this margin.

**Figure 7:**
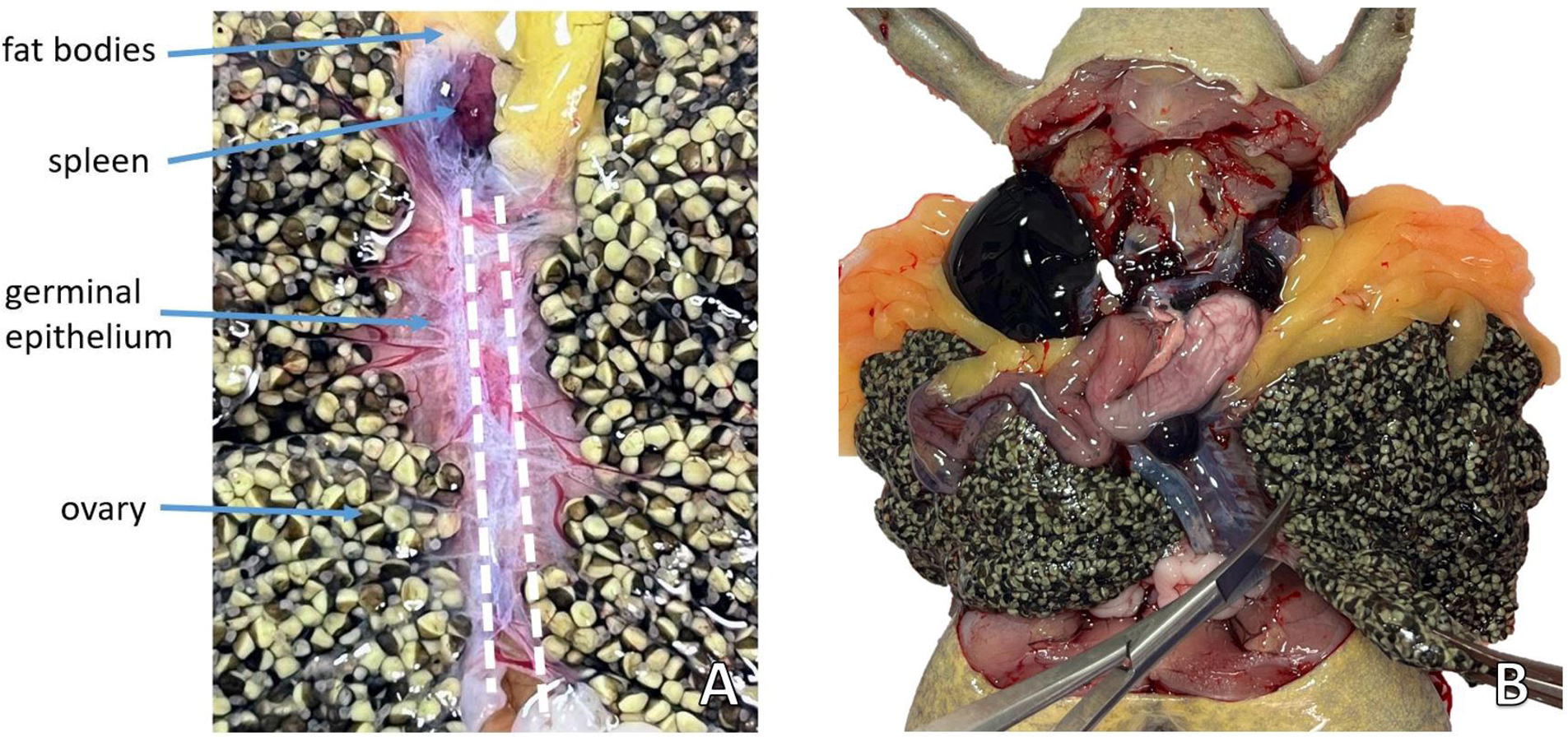
**Ovary removal of an unperfused *X. laevis.*** (**A**) With the ovary lobes on their respective sides, the continuity of the germinal epithelium to the peritoneal wall (over the kidneys) is visible. Two white dashed lines indicate where to sever these attachments. (**B**) An ovary is pulled away from the paired kidneys and cut where the germinal epithelium meets the peritoneum.

**Figure 8:**
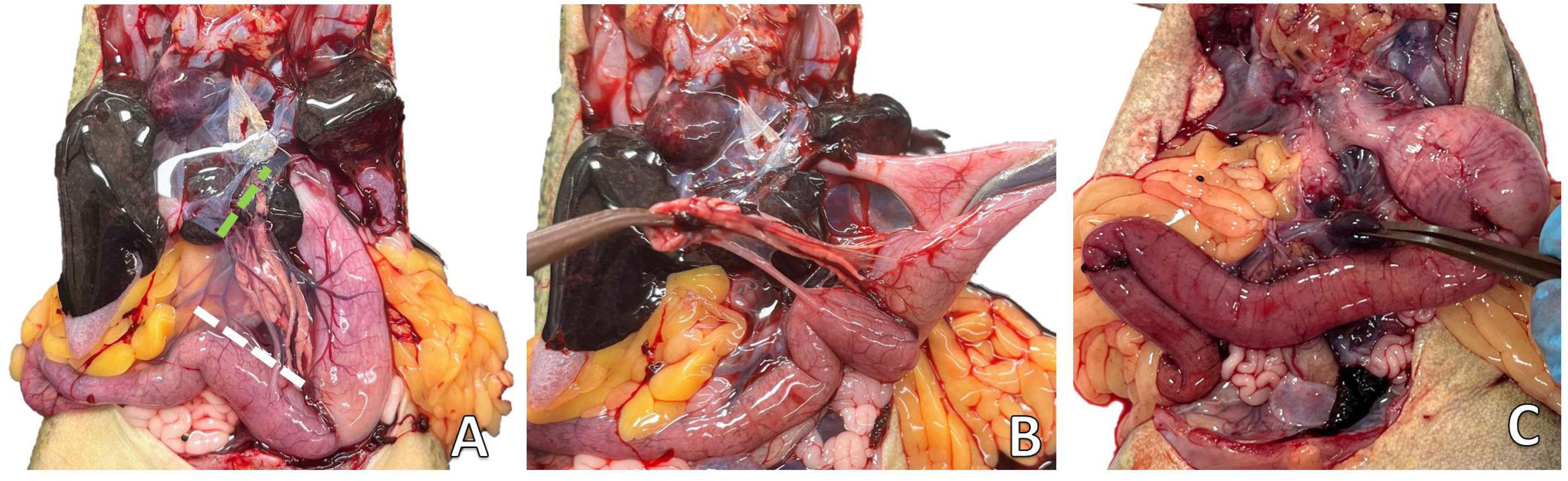
**Pancreas and spleen sampling of an unperfused *X. laevis.*** (**A**) The pancreas with the hepatopancreatic duct and ligament visible. A green dashed line indicates where to sever the pancreas from the anterior lobe of the liver. A white dashed line indicates where the mesentery may be cut to separate the pancreas from the duodenum. (**B**) The pancreas is teased off of the stomach. (**C**) The spleen is sampled at the superior end of the kidneys.

**Figure 9:**
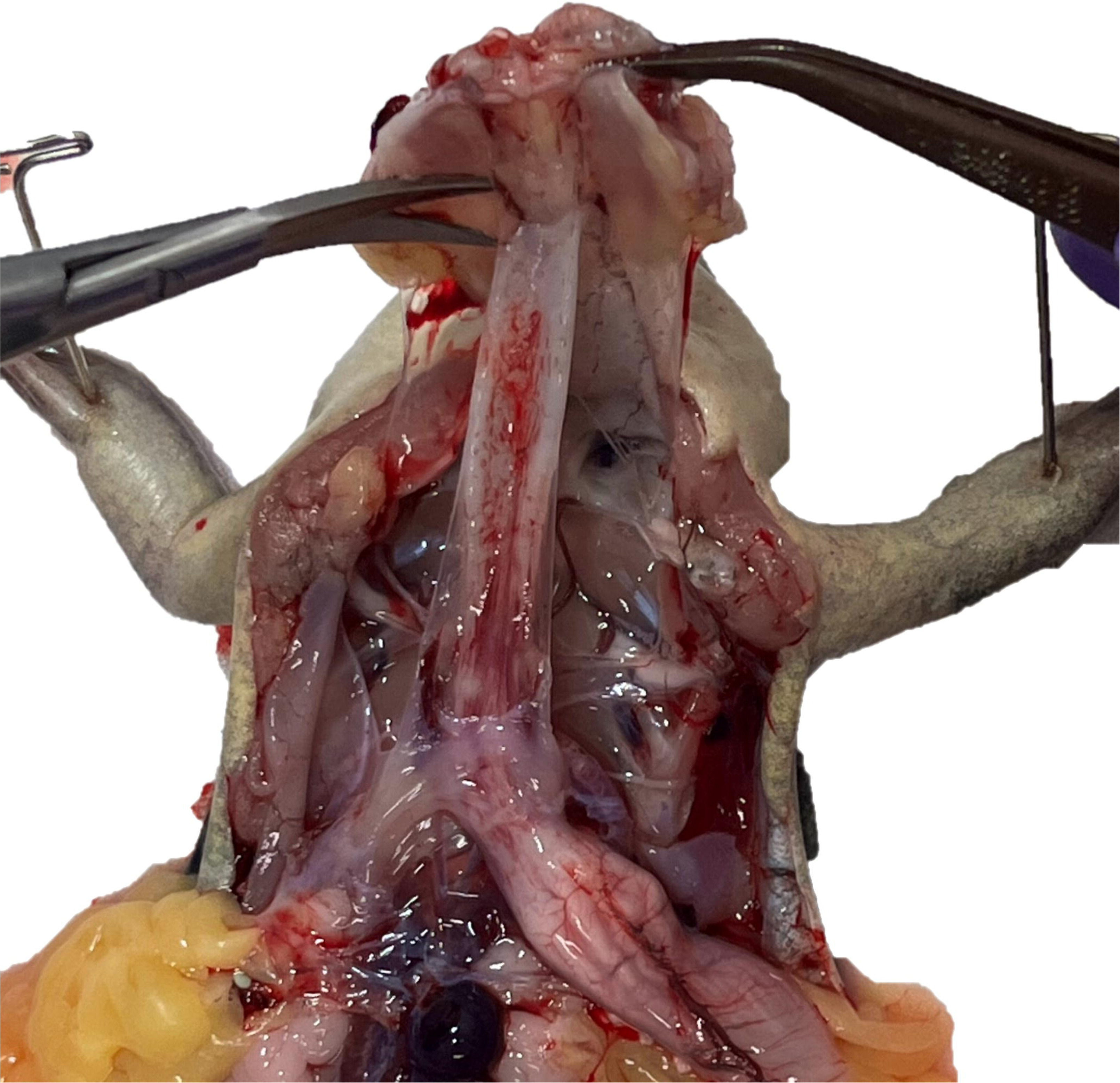
**Larynx removal of an unperfused *X. laevis.*** The larynx has had the bronchus of each lung removed and has been severed from its inferior clear attachments. The larynx is now reflected out of the cavity and methodically cut away from the esophagus.

**Figure 10:**
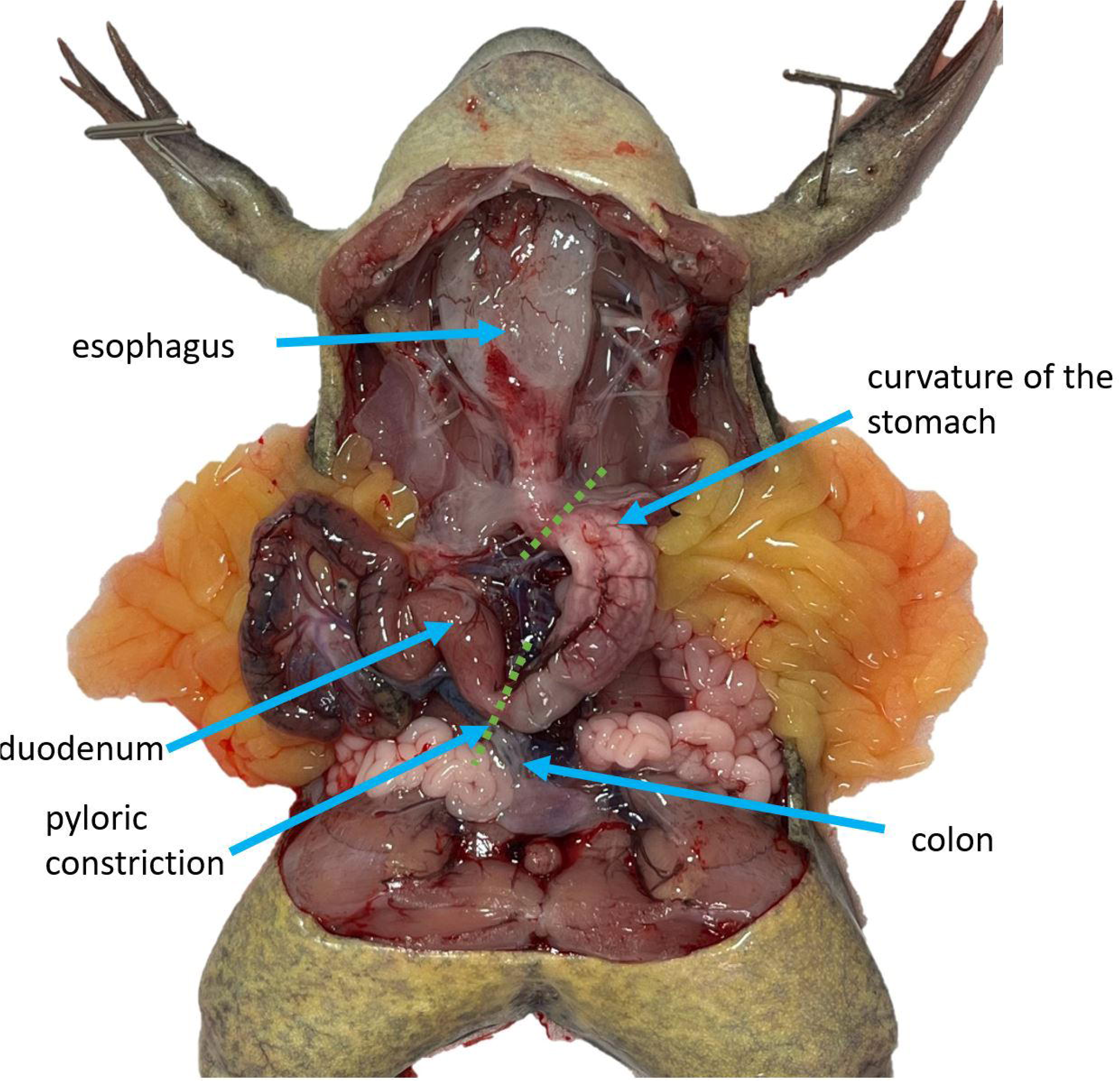
**Alimentary canal of an unperfused *X. laevis.*** With the previous samples as well as the ovary having been removed, the alimentary canal is now fully visible including the esophagus, the curvature of the stomach, the pyloric constriction, the duodenum, and the colon. Green dashed lines indicate the margins on either end of the stomach.

**Figure 11:**
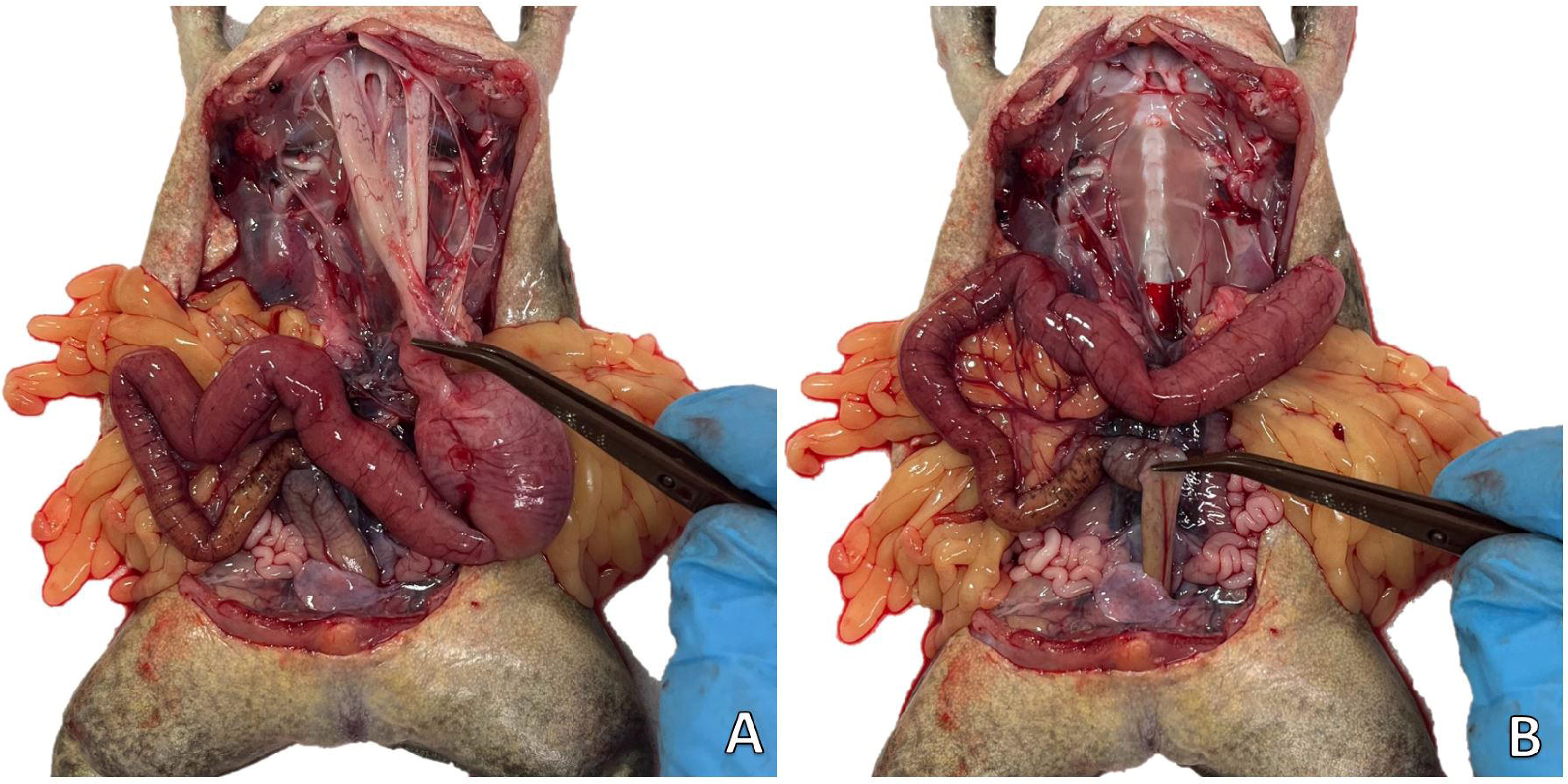
**Esophagus and intestine detachment of an unperfused *X. laevis.*** (**A**) The esophagus can be easily severed from the buccal cavity when pulled taut. Note a section has been removed to make the Eustacean opening visible. (**B**) The colon is pulled taut to be severed close to the cloaca.

**Figure 12:**
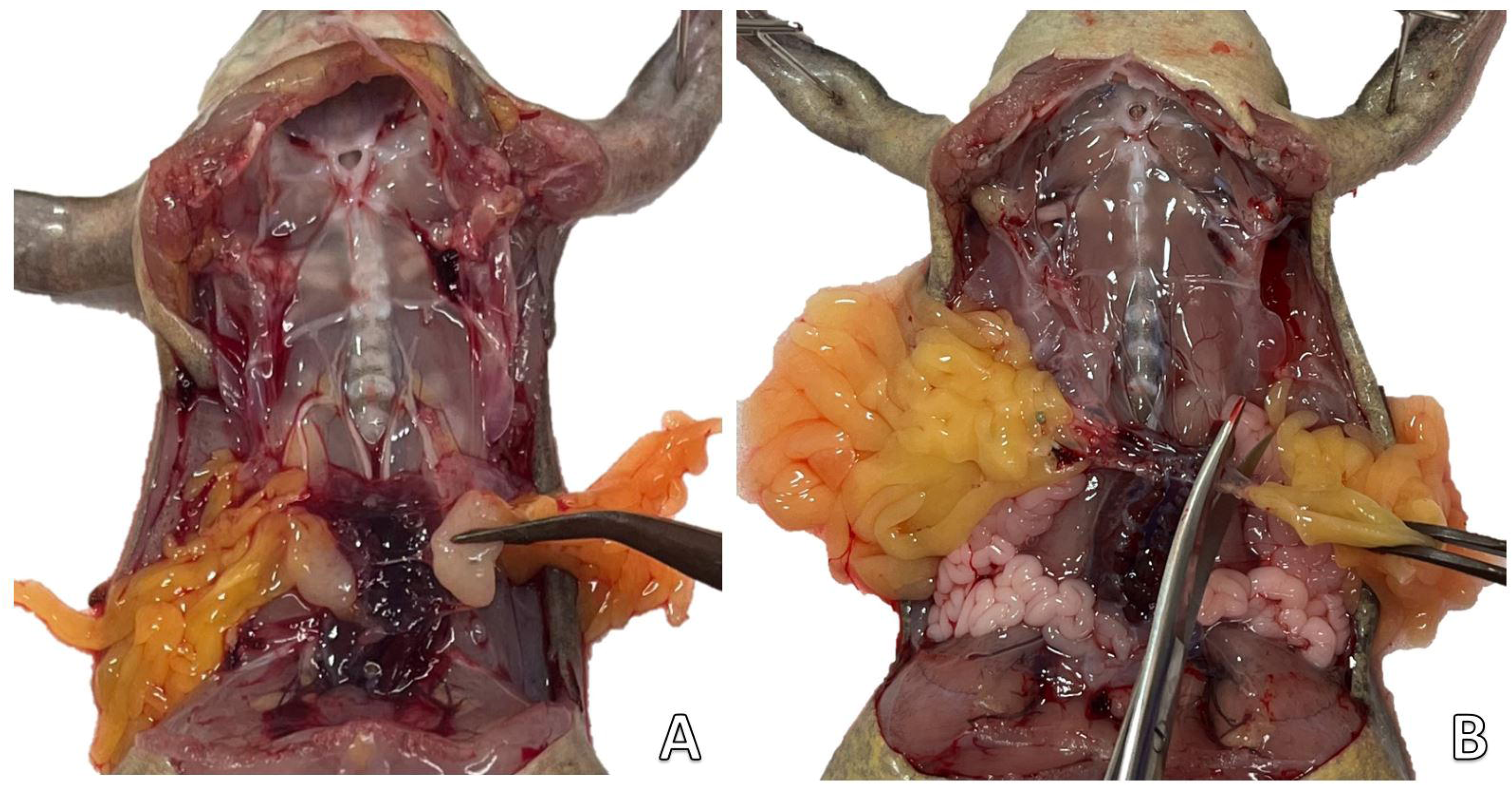
**Testes and fat body sampling of an unperfused *X. laevis.*** (**A**) In males, testes are identified over the kidneys and cut away from the peritoneum. (**B**) By pulling at the mass of the fat body its attachment to the peritoneum, over the kidney, is visible. Cut it away from the kidney leaving a margin.

**Figure 13:**
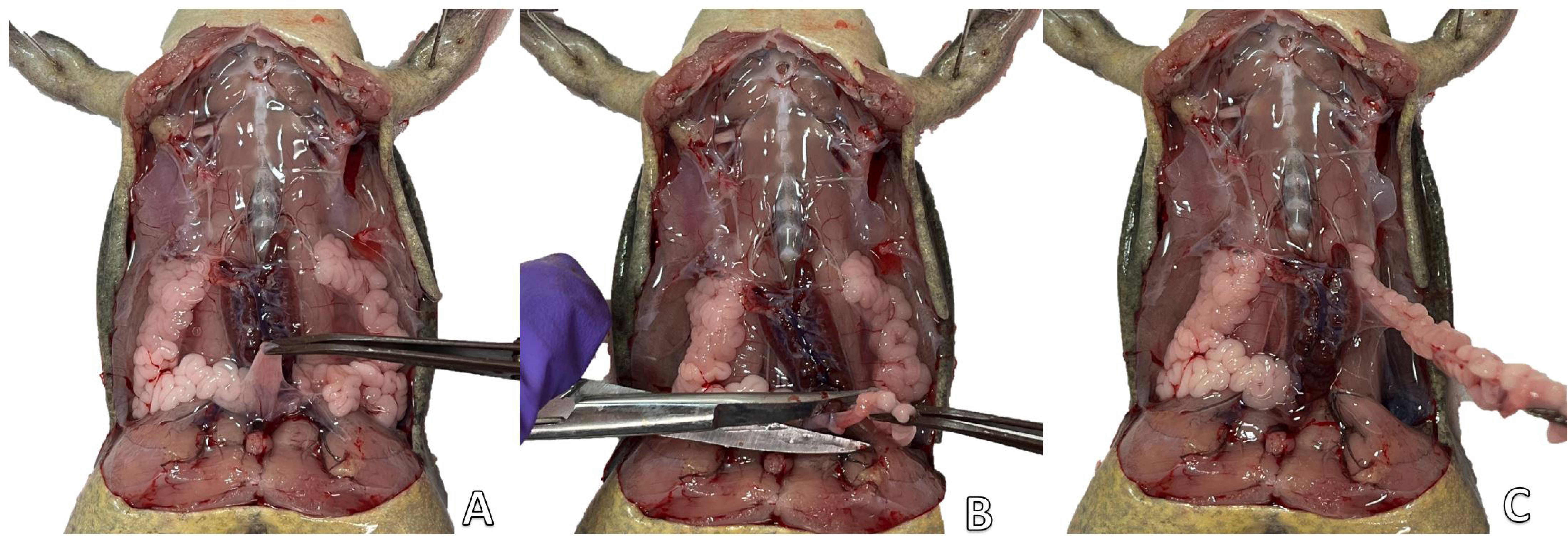
Urinary bladder and oviduct sampling of an unperfused female *X. laevis* . (**A**) The urinary bladder is tugged upward so that it can be severed as close to the cloaca as possible. (**B**) the oviduct is drawn away from the cloaca so that it too may be cut as close to the cloaca as possible. (**C**) The oviduct is lifted out of the coelomic cavity. Clear attachments to the kidney become apparent and should be severed.

**Figure 14:**
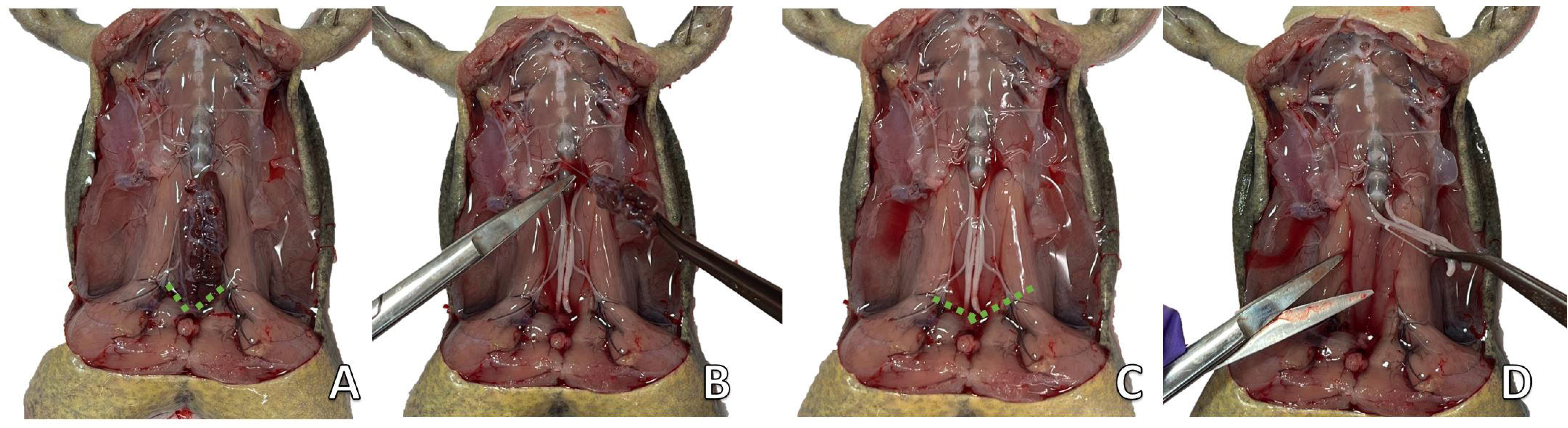
**Kidney and sciatic plexus sampling of an unperfused *X. laevis.*** (**A**) With all the previous tissues removed the kidney is fully visible, green dashed lines indicate where to sever the peritoneum of the kidney, at its inferior end. (**B**) The kidneys’ remaining peritoneal attachments are cut as the kidney is lifted out of the coelomic cavity. (**C**) The sciatic plexus is now fully visible. Green dashed lines indicate where the sciatic plexus’s nerve bundles should be cut where they exit the coelomic cavity. (**D**) The sciatic plexus is then lifted out of the coelomic cavity and severed where it originates from the spinal canal.

**Figure 15:**
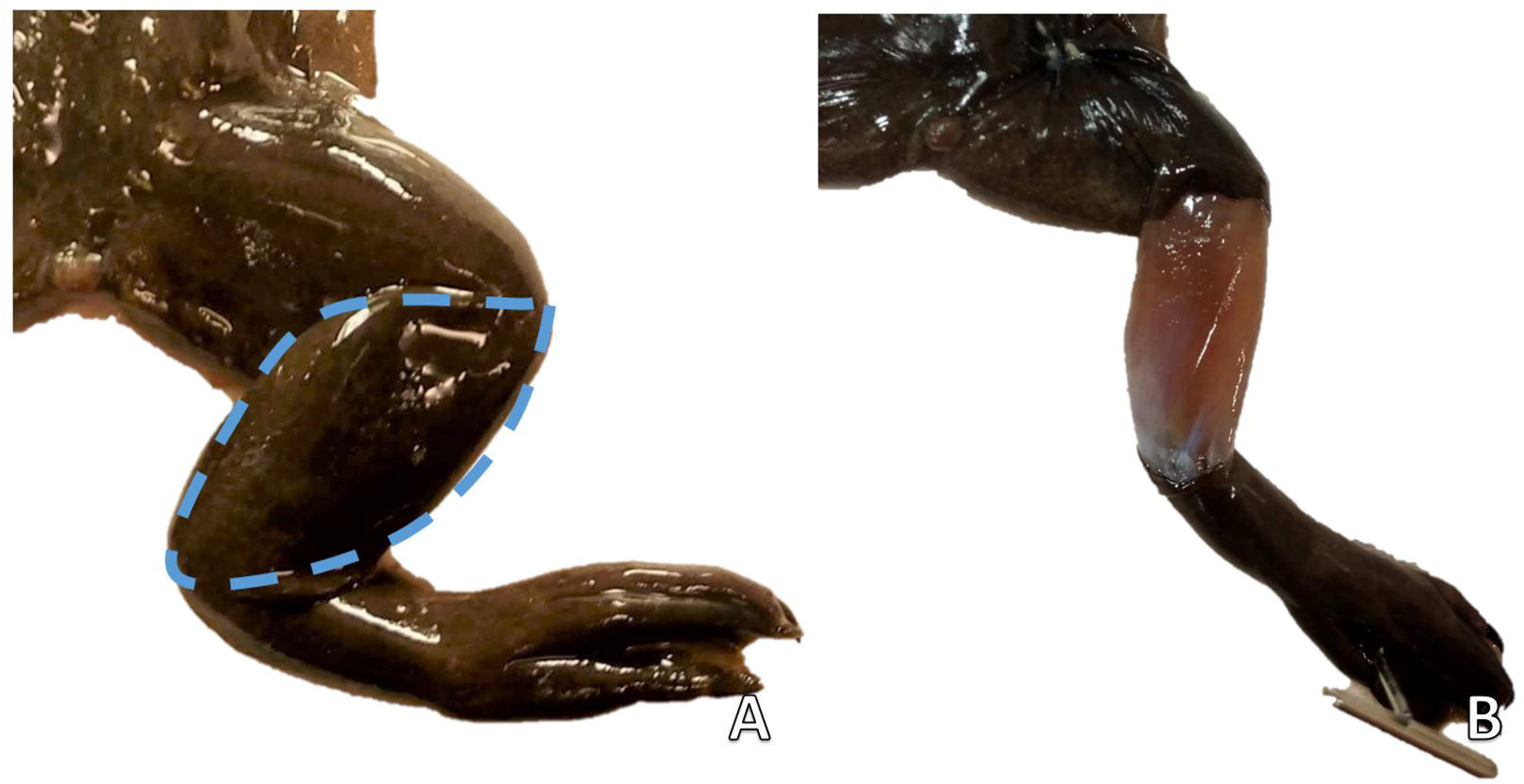
**The skin sampling of an unperfused *X. tropicalis.*** (**A**) The right leg with a dashed line indicating the area of skin to be sampled. (**B**) The right leg with a skin sample removed over the tibiofibula.

**Figure 16:**
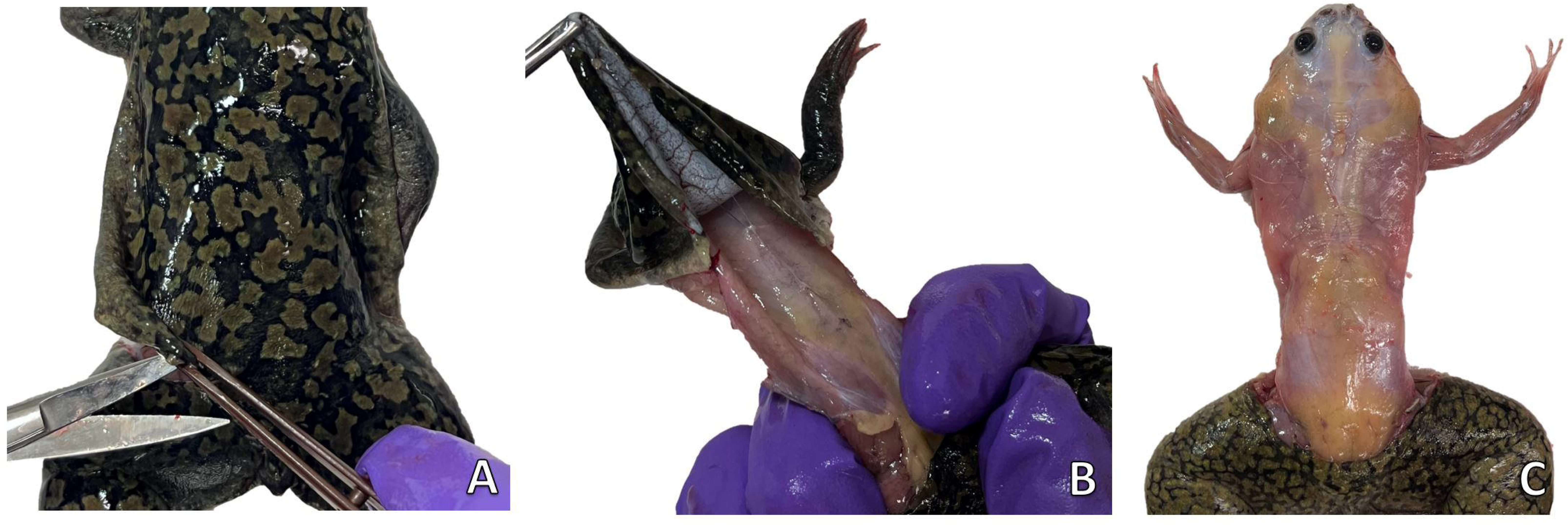
The superior skinning of an unperfused *X. laevis* . (**A**) The dorsal skin is cut laterally so that there is no attachment between the skin of the superior and inferior body. (**B**) The dorsal skin is being pulled forward, over the head, skinning the body. (**C**) An unperfused *X. laevis,*once the superior portion of its body has been skinned

**Figure 17:**
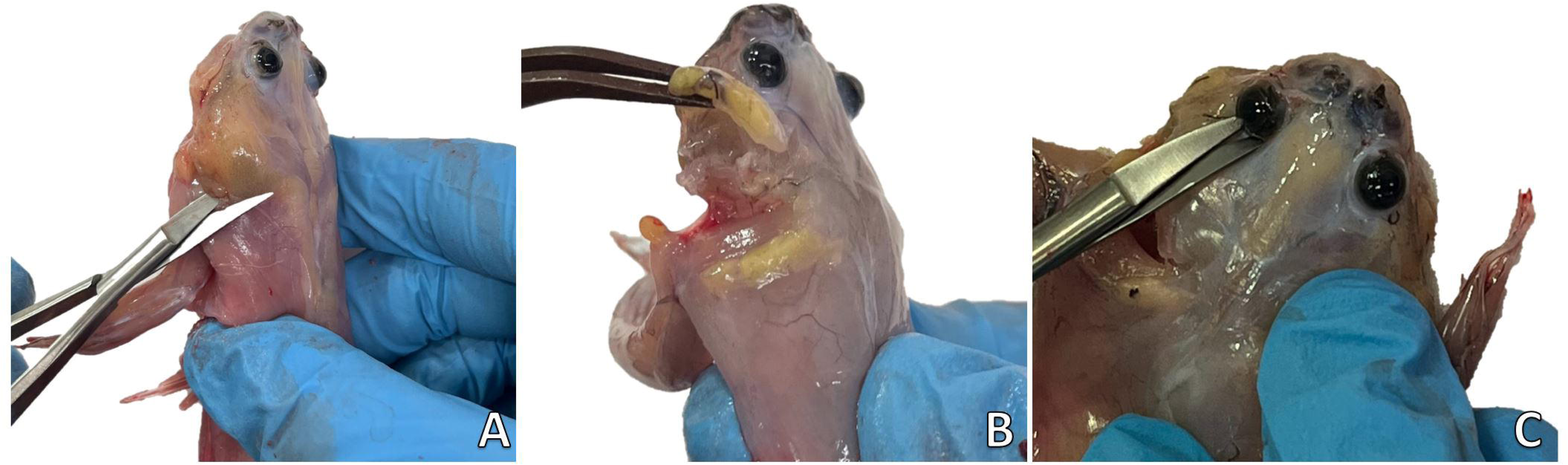
**Thymus and eye sampling of an unperfused *X. laevis.*** (**A**) The thymus and associated fatty mass are cut away from the jaw, at the posterior end. (**B**) The mass is then reflected away from the body revealing a cluster of melanophores, that is the thymus. Note that the cluster of melanophores will vary drastically depending on the animal’s maturity. (**C**) Curved iridectomy scissors are inserted around the eye. They are then used to sever the orbital muscles and optic nerve.

### Representative results

By utilizing **Figure 1** to **Figure 17** and following all steps of this protocol the heart ventricle, arterial trunk, left liver lobe, gallbladder, lung, pancreas, spleen, larynx, esophagus, stomach, intestines, testes, fat bodies, oviduct, paired kidneys, sciatic plexus, skin, thymus, and whole eye were cleanly excised within an hour of euthanasia. Within this time, the samples are rinsed and trimmed so that they will appear, as shown in **Figures 18 and 19**.

**Figure 18:**
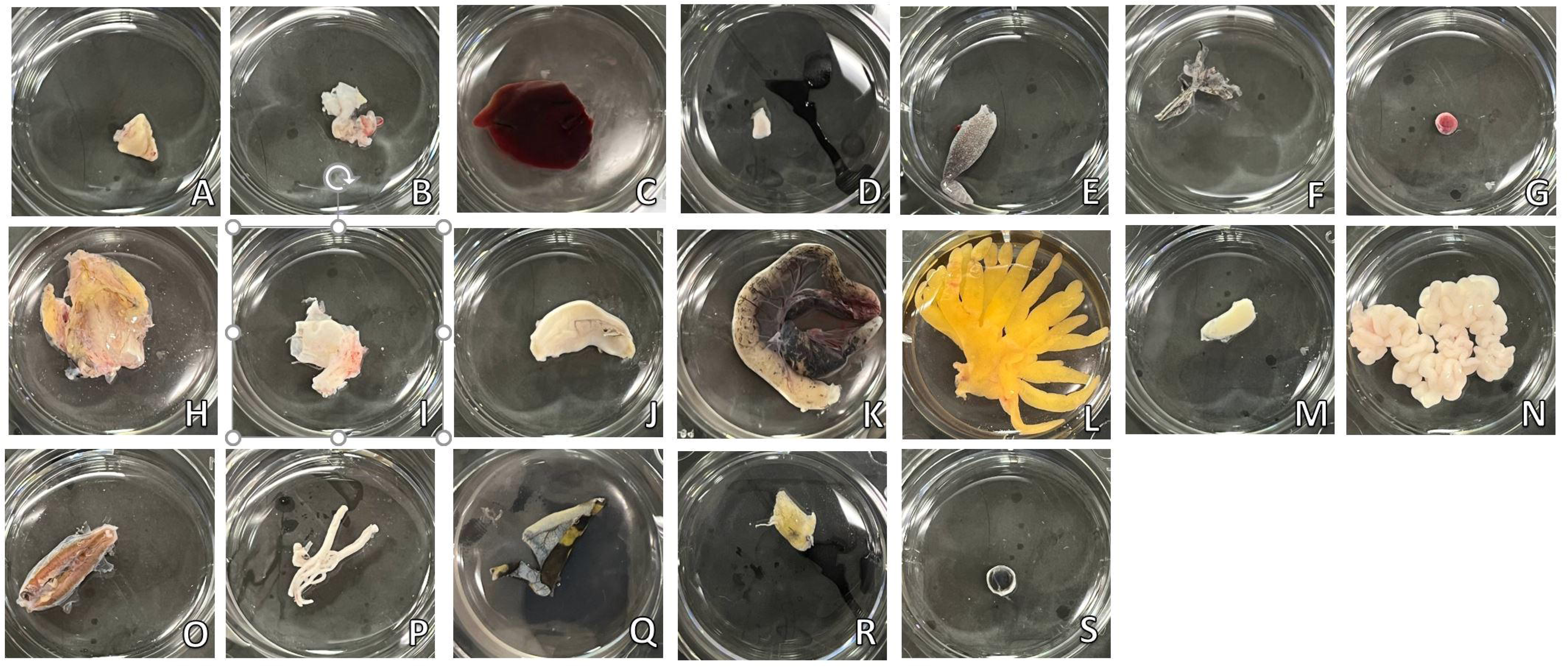
**Representative results: perfused samples of an *X. laevis.*** All samples are imaged in the wells of a six-well plate. (**A**) heart ventricle, (**B**) arterial trunk, (**C**) liver, (**D**) gallbladder, (**E**) lung, (**F**) pancreas, (**G**) spleen, (**H**) larynx, (**I**) esophagus, (**J**) stomach, (**K**) intestines, (**L**) fat bodies, (M) testis, (**N**) oviduct, (**O**) paired kidney, (**P**) sciatic plexus, (**Q**) skin, (**R**) thymus and (**S**) eye.

**Figure 19:**
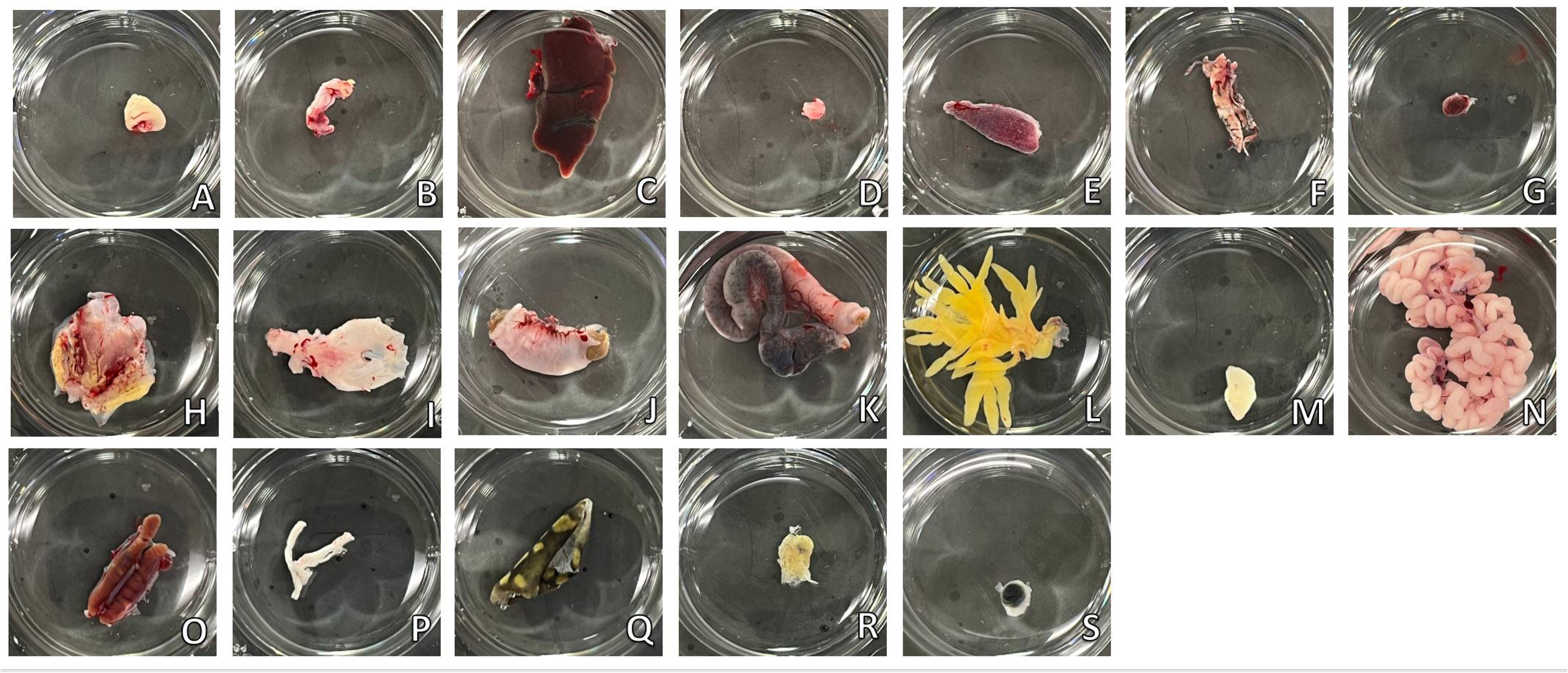
**Representative results**: **unperfused samples of *X. laevis*** . All samples are imaged in the wells of a six-well plate. (**A**) heart ventricle, (**B**) arterial trunk, (**C**) liver, (**D**) gallbladder, (**E**) lung, (**F**) pancreas, (**G**) spleen, (**H**) larynx, (**I**) esophagus, (**J**) stomach, (**K**) intestines, (**L**) fat bodies, (**M**) testis, (**N**) oviduct, (**O**) paired kidney, (**P**) sciatic plexus, (**Q**) skin, (**R**) thymus and (**S**) eye.

## Discussion

The protocol presented here is designed to provide flexibility for users at various experience levels while ensuring the collection of essential tissues in a timely and efficient manner. New users are advised to follow each step in the protocol as outlined to ensure the systematic collection of tissues. More experienced users, however, may opt to skip certain steps and sample only the tissues they require. For instance, accessing the spleen, which is located behind the mesentery, can be achieved without sampling the pancreas by simply shifting the stomach. This adaptability allows for customization of the sampling process according to specific research needs.

While the protocol is designed for efficiency, enabling the collection of up to 18 tissues in under an hour, further refinement of techniques may be necessary for specific applications. For example, users may choose to sample the mesentery separately from the intestines or collect the adrenal glands independently from the kidneys. Additionally, the gastrointestinal organs could be flushed with extra care to preserve their integrity^12^. These adjustments could enhance the accuracy and completeness of the data, particularly when dealing with highly specialized tissues.

The method was also developed to be compatible with rapid perfusion^8^, a crucial step for various downstream applications such as proteomics, where tissue freshness is also critical. This protocol represents part one of a comprehensive tissue sampling guide, which will include additional tissues not described in this method. For users seeking a more concise approach, an abridged version of the protocol is available for sampling a select range of tissues, including the heart ventricle, liver lobe, pancreas, fat bodies, paired kidneys, and skin^13^.

Although this method was initially developed for use in *Xenopu*,*s*it is versatile enough to be applied to other herpetological models with minimal modifications^14^. Furthermore, the protocol is designed to be compatible with both male and female specimens, ensuring broad applicability across different species and research objectives.

Future iterations of this method may focus on refining tissue-specific techniques to further enhance the protocol’s utility in a variety of biological research contexts.

## Supporting information

Materials list

## ACKNOWLEDGMENTS

This work was supported by NIH’s OD R24 grant ODO31956. We would also like to thank Samantha Jalbert and Jill Ralston for their assistance and support as well as our editor and anonymous peer reviewers for their feedback.

