## Supplementary material for "Efficient Techniques for Comprehensive Tissue Sampling in Adult Xenopus": Materials list

| Name of Material/ Equipment | Company | Catalog Number | Comments/Description |
|---|---|---|---|
| 5X Magnifying Glass with LED Light and Stand | amazon.com | B08QJ6J8P1 | light must not produce heat |
| Disposable Transfer Pipets | VWR | *414004-036* | |
| Dissecting Fine-Pointed Forceps | Fisher Scinetific | 08-875 | |
| Dissecting scissors sharp piont, straight 6.5" | VWR | 76457-374 | |
| Dissection Tray | Fisher Scinetific | 14-370-284 | styrofoam sheets are an acceptable alternative |
| Euthanasia container | US Plastic | Item 2860 | alternative opaque containers acceptable |
| Euthanasia container lid | US Plastic | Item 3047 | |
| Iridectomy Scissors 6" | VWR | 470018-938 | iris scissors are an acceptable alternative |
| MS-222: Syncaine (formerly tricaine) | Pentair AES | TRS1 | |
| PBS x 1 | Corning | 21-040-CV | |
| Rat Tooth Tissue Forceps 5.5 in., Stainless Steel | Fisher Scientific | S08100 | |
| Sodium Bicarbonate, Powder, USP | Fisher Scientific | 18-606-333 | |
| Specimen Forceps, Serrated | VWR | 82027-442 | |
| T-Pins for Dissecting | Fisher Scinetific | S99385 | |
